# Electrophysiological correlates of associative recognition memory for identity, spatial, and temporal relations

**DOI:** 10.1101/2022.10.22.513054

**Authors:** Ofer Hugeri, Eli Vakil, Daniel A. Levy

## Abstract

The formation of associative representations and their retrieval from episodic memory are vital cognitive functions. However, it is unclear to what extent retrieval of the basic component relations of episodic memory – identity, time, and space – requires different or shared brain mechanisms. In the current study, we employed EEG to track the time courses of electrophysiological correlates of retrieval processes of memory for identity relations, temporal order, and spatial configuration. Participants engaged in pair-associate learning of serially presented and spatially configured object picture pairs, followed by discrimination of identity, spatially, or temporally intact and rearranged pairs. Event-related brain potentials (ERPs) revealed distinct patterns of activity during successful retrieval of identity, spatial, and temporal relations that differed by the status of association, across the three retrieval time windows examined (300-500, 500-800, and 800-1000 ms). The identity relations condition was distinguished by a widespread greater negative-going deflection for rearranged relative to intact pairs in all three time windows. For the temporal relations condition, we observed a widespread more negative-going deflection for rearranged than intact pairs, significant in the second time window only. For the spatial relations condition, there was a widespread positive-going deflection greater for rearranged than for intact pairs, significant in the early and in middle time windows. These patterns of activity suggest that retrieval of associative memory traces for identity, spatial, and temporal relationships involve dynamically different processes, which may partially rely on different sets of basic associative mechanisms.

## 1. Introduction

A key characteristic of episodic memory, and what distinguishes it from semantic memory, is the conscious remembrance of events in which people or things are situated in time and space (Tulving, 1972). The retrieval of episodic associations -that two or more stimuli were experienced in close proximity -from long-term episodic memory is a crucial cognitive function. It enables us to reconstruct environments in which we have been present, sequentially order events in which we have participated, and relive incidents that we have experienced. Episodic associative memory rests on three main pillars: the ability to remember which combination of people or objects participated in a given event (item-identity associative memory), to remember the relative whereabouts of those items (associative spatial location memory), and to retrieve the sequence in which the items appeared relative to each other (temporal order memory) (Tulving, 1983). While associative memory may be reflected in the recollective aspects of item recognition (e.g., ‘remember’ responses in the remember-know paradigm and various source judgments; Mitchell & Johnson, 2009; Yonelinas, 2002) it is expressed most directly in examination of cued recall and associative recognition. The latter is the focus of the current study, as it provides an effective assay of the spatio-temporal aspects of associative memory, as described below.

Progress in understanding episodic memory may be achieved by exploring the extent to which these three components of associative memory converge or diverge in their processes and brain mechanisms. Numerous studies have highlighted the hippocampus, the prefrontal cortex, and their interactions as shared fundamental bases for these episodic memory abilities (see Eichenbaum, 2017, for a review). Several studies have identified the significant role of the hippocampus in remembering events in the spatial and temporal context in which they occurred, using human and non-human models (e.g., Butterly et al., 2012; Eichenbaum, 2004; 2017). Barker et al. (2017) and Chao et al. (2016) have shown that for remembering where and when stimuli were presented hippocampal-medial prefrontal cortex interaction is crucial, to which Chao et al. (2020) added the contribution of lateral entorhinal cortex. On the other hand, several studies examining cognitive and behavioral (e.g., Tolentino et al., 2012; Van Asselen et al., 2006) and physiological (e.g., Ekstrom & Bookheimer, 2007; Staresina & Davachi, 2009) indices of memory for temporal order, relative spatial location, and inter-item (identity) associations suggest that these aspects of episodic memory might be based on partially different brain mechanisms.

A study by Rajah et al. (2011) tracked patterns of prefrontal cortex activity during spatial and temporal context memory retrieval. In that study, participants viewed pictures of unfamiliar faces presented serially and across three possible screen locations, and were subsequently asked to arrange the face pictures in either spatial or temporal original orders. Analyses of fMRI data focusing on prefrontal cortex generally found a similar patterns of activation for both tasks. In another study, Rajah et al. (2010) had participants watch a series of video episodes, and subsequently choose either the scene that happened earlier (temporal order judgment) or the scene with a correct object spatial arrangement (spatial location judgment). In that paradigm, fMRI indicated that the precuneus and angular gyrus were associated with temporal retrieval, while a dorsal fronto-parietal network was engaged during spatial retrieval. A similar task was employed by Kwok et al. (2012), who report similar patterns of activation. However, as noted by Rajah et al. (2011), in that paradigm task structure and experimental design differed for temporal and spatial context tasks, and performance was not equated for accuracy between tasks. These findings thus provide mixed support for process-specific dissociation of retrieval.

Another line of research addresses the issue by examining possibly differential effects of healthy aging on associative memory for identity, spatial, and temporal relations. Some studies have indicated that memory abilities for temporal order, spatial location, and inter-item associations may differ across the lifespan (Benjamin, 2016; Bridger et al., 2017; Old & Naveh-Benjamin, 2008; Ratcliff et al., 2015). However, such findings are not uniform. For example, while Diamond et al. (2018) found spared temporal and spatial memory, and Cheke (2016) reported impaired temporal but spared spatial memory, other studies reported impairment of either temporal associative memory (Cabeza et al., 2000; Fabiani & Friedman, 1997; Kausler et al., 1990; Parkin et al., 1995; Vakil et al., 1997), spatial associative memory (Lemay & Proteau, 2003; Kessels et al., 2005), or both (Kessels et al., 2007; Vakil & Tweedy, 1994). Moreover, many studies have shown an impairment in inter-item memory in older adults (Badham & Maylor, 2013; Bridger et al., 2017; De Brigard et al., 2020; Giovanello & Schacter, 2012; Greene & Naveh-Benjamin, 2020).

In an attempt to resolve these divergent findings, we designed a paradigm in which basic associative recognition memory for identity, spatial configuration, and temporal order relationships between pairs of visual objects is tested via discrimination of intact and rearranged pairs (Hugeri et al., 2022). The paradigm is fully described below, but for now we note that it is based on the approach that the most basic building blocks of episodic memory are the identity, temporal and spatial relationships between pairs of items within a particular context. It differs from those used in prior research by examining all three associative aspects within the same framework, holding stimuli and presentation constant, with the only difference between conditions being the type of associative relationship which the participant needs to remember. Using this paradigm, we previously examined the extent to which accuracy and response time in each element is affected by healthy aging. Age-related declines in associative memory were observed equally for all types of associations, but these declines differed by associative status: aging most strongly affected the ability to discriminate rearranged pairs. These results suggest that associative memory for identity, spatial, and temporal relationships are equally affected by healthy aging, and may all depend on a shared set of basic associative mechanisms

To determine whether the consonance between temporal, spatial, and identity associative processes suggested by aging effects is characteristic of brain mechanisms supporting associative memory, anatomical and physiological investigation is required. The current study explored the processes underlying retrieval of identity, temporal and spatial relations by examining event related potential (ERP) correlates of retrieval in an episodic “minimal pairs” paradigm. While lacking the ability to anatomically identify the brain regions or networks supporting the three aspects of associative episodic recognition, the electrophysiological correlates of the associative retrieval processes do provide high-resolution data about their time courses. Additionally, it is possible to examine how the patterns of EEG activity associated with these retrieval types relate to well-documented ERP correlates of item-associative recognition processes (e.g., Donaldson & Rugg, 1998; Jäger, Mecklinger, & Kipp, 2006). Retrieval of associative memory is considered to employ recollective processes involving episodic reconstruction of encoding context. However, in certain circumstances, it can also be accessed by familiarity processes, when encoding conditions enable the formation of a unitized representation (Quamme et al., 2007; Tibon et al., 2014; for review, see Mecklinger & Jäger, 2009; Yonelinas et al., 2010). We hypothesized that this could be the case for identity relations, and therefore predicted that successful identity associative recognition might be indexed by a response akin to the familiarity-related FN400 ERP component, but that this would not be the case for the spatial and temporal associations. In contrast, in later time windows in which ERP components are associated with recollective processes (Curran & Rugg, 2007), it is possible that responses evoked during spatial and temporal associative recognition might be more prominent. Exploring the differences between the time course of basic retrieval processes for temporal, spatial and identity associative relations will provide a fuller picture of associative recognition memory mechanisms.

## 2. Methods

### 2.1 Participants

Thirty-three healthy young adults participated in the experiment. All participants were right-handed (positively scored on the Edinburgh Handedness Inventory; Oldfield, 1997), with normal or adjusted-to-normal vision, self-reportedly with no psychiatric or neurological disorders, seizures, or use of psychotropic substances. Four participants were excluded from the analysis: two due to technical failure, one due to incompatibility with the screening criteria which was revealed retrospectively, and one due to withdrawal during the experiment, leaving 29 participants (18 females, mean age = 23.8, *SD* = 2.7, range 19-31) whose data was analyzed. Participants were compensated in the form of academic requirement credit or payment and provided written informed consent for a protocol approved by the human participants research ethics committee of Reichman University.

### 2.2 Stimuli

The experimental stimuli comprised a set of 96 common object pictures (e.g., wheelbarrow, crib, cucumber, cat, pizza, paper clip) drawn from the Bank of Standardized Stimuli (BOSS) (Brodeur et al., 2014). All images employed were rated highly nameable by an independent panel of participants who did not take part in the main study. Each image was resized to 135 × 135 pixels to reduce eye movement during stimulus presentation, and edited to have an all-white background. Stimuli were assigned to three sets of 32 pairs of semantically unrelated pictures; pair stimuli unrelatedness was confirmed by independent raters who did not take part in the main study. These three sets were assigned to identity, spatial, and temporal task conditions, counterbalanced across participants. Additional pictures were added and paired as required for examples and practice trials, as described below.

### 2.3 Procedures

#### 2.3.1 Task structure

To track the electrophysiological correlates of identity-spatial-temporal associative retrieval processes and their time courses, we strove to construct a paradigm in which the tasks assessing those forms of associative memory would be as closely matched as possible. The basic task in all cases was to intentionally form associative memories for a set of several pairs of object pictures using a deep encoding task, and thereafter to make confidence-scaled recognition judgments on a set of probe pairs, each of which was either identical to a studied pair or rearranged in the fashion relevant to the type of memory being assessed.

At the beginning of each task, participants were informed that they were about to see a pair of object pictures on the screen, memory for which would later be tested. They were told that each picture would appear twice, and that each time a different picture would be presented with it. At encoding, in each trial, one picture was presented above or below fixation for a certain exposure time (detailed in Table 1), followed immediately by a second picture in the opposite location (as depicted in Figure 1). For the deep encoding procedure, participants were instructed to make an association for each pair that related to the aspect of the associative relationship (identity, spatial, or temporal) being tested in that part of the experiment. They were asked to make the association as vivid as possible in order to remember the pictures’ relationship. So, for example, in the temporal condition, if the participants saw a picture of a candle and then a shoe, they were to think about themselves first lighting a candle and then proceeding to use the light to look for their shoe. The experimenter demonstrated the first practice trial, after which the participant performed four rounds of practice aloud while receiving feedback on the associations. After the practice phase, all associations were made silently. Importantly, in each encoding block, each constituent picture was used twice, to construct two associative pairs, with a different associate in each pair. A single aspect of the identity, spatial, or temporal characteristics of each picture differed between the two pairs in which it appeared. For example, when paired with a car, a picture of a dog might appear first and above fixation, while in its second appearance, paired with a banana, it might appear second and above fixation (in blocks testing temporal associative memory), or first and below fixation (in blocks testing spatial associative memory), or it might appear in the same place and order (in blocks testing identity associative memory) (see Supplementary Table 1). The effect of this double-pairing was to require encoding and retrieval of spatio-temporal and identity information that was specific to each associative pairing, such that neither single-item location, nor single-item temporal order information, nor memory for a single identity association would enable successful subsequent retrieval.

**Table 1.**
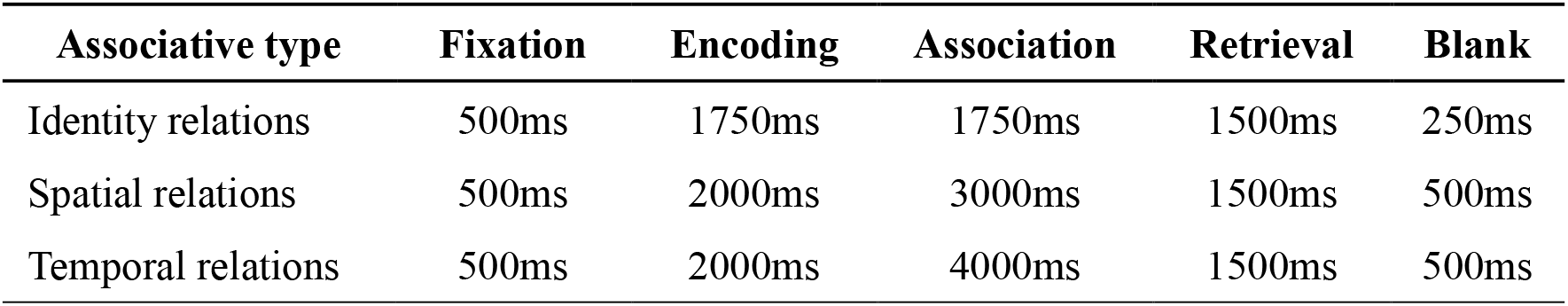
Stimulus display time (ms) for the Identity, Spatial, and Temporal relations.

**Figure 1.**
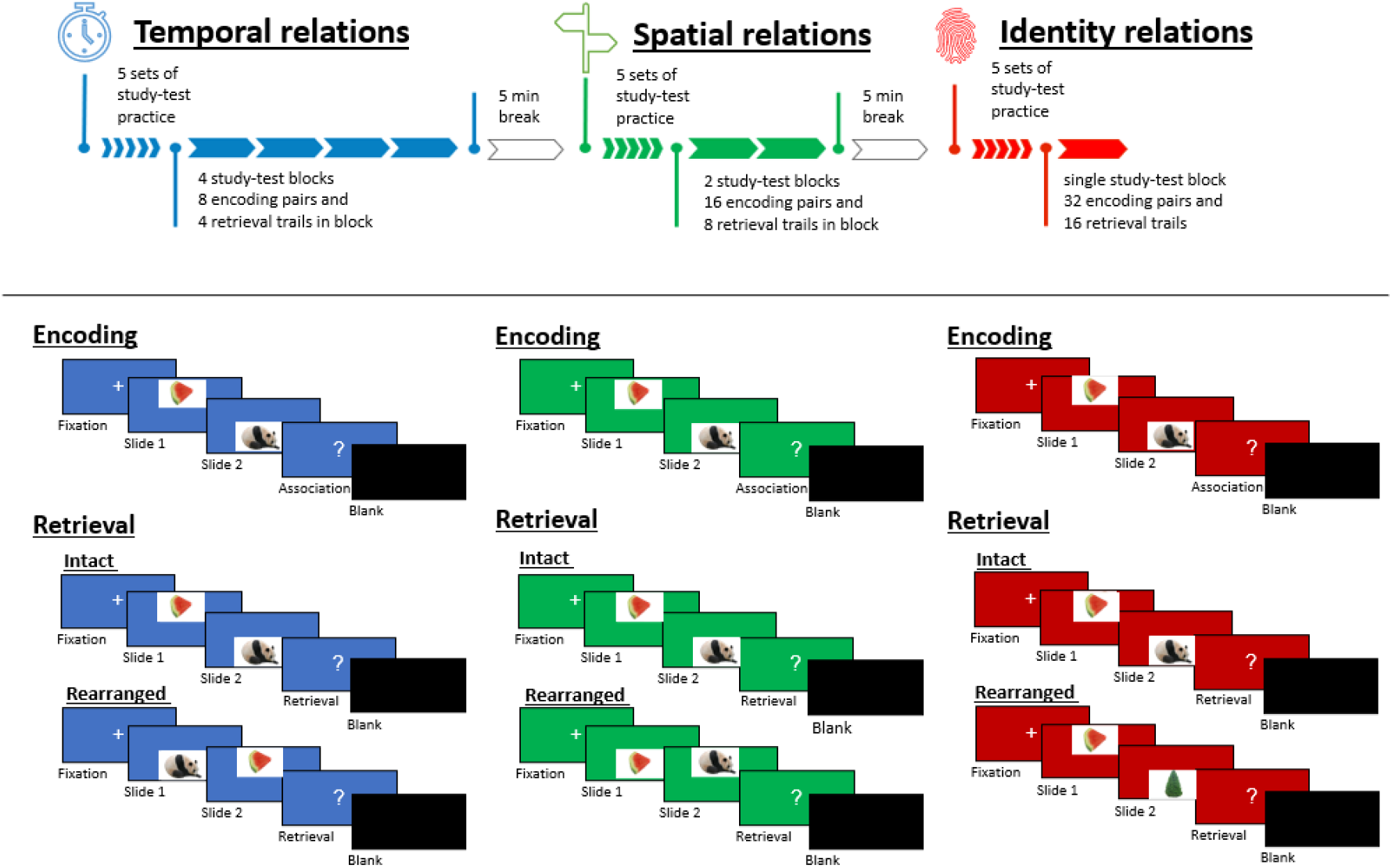
Experimental design. Participants performed three minimal pair associative recognition tasks for temporal, spatial and identity relations. In each task, participants learned to associate two consecutively presented object pictures and were instructed to focus only on the task-relevant dimension (order, location, or identity). Every picture appears twice in each encoding block, each time paired with a different picture.

A test block immediately followed each encoding block. At test, each probe pair was presented in the same format as at encoding (i.e., serial presentation of pictures in different locations on the screen) and was either identical to a studied pair in all dimensions, or differed in a single dimension (identity, spatial, or temporal), in accordance with the type of memory being assessed in the relevant block. Thus, a rearranged pair in the trials assessing associative memory for spatial relations would display the same two pictures as at encoding, each appearing in the same serial order position as at encoding, but with the locations of the pictures switched. A rearranged pair in the trials assessing associative memory for temporal relations would display the same two pictures as in encoding, each appearing in the same spatial position as at encoding, but with the order of appearance of the pictures switched. A rearranged pair in the trials assessing associative identity memory would display two pictures that were not paired at encoding, but with each appearing in the same spatial position and temporal order as it did in an encoding trial. Importantly, at test, each item was only presented once, in one of the two configurations used at the study. Thus, none of the associative recognition judgments could be informed by decisions made on earlier trials.

#### 2.3.2 Procedure

The experiment included two sessions which took place on different days (mean spacing = 6.7 days, *SD* = 1.6). This was required to avoid participant fatigue during collection of the large number of experimental trials required to insure sufficient EEG artefact-free trials in all six test conditions. Each of the sessions lasted approximately 1.5 hours. Participants were tested individually in a quiet room. After providing informed consent, they filled out the Edinburgh Handedness Inventory (Oldfield, 1971). They were seated at a distance of ∼70 cm from a 24-inch computer monitor with a 144-Hz refresh rate (AOC Freesync G2460PF). Following EEG electrode cap preparation (described below), participants received instructions explaining the procedures to be followed during the study and test stages of the experiment, as described above. Before the presentation of the first study block in each part of the experiment, participants were explicitly instructed to focus only on the relevant dimension of that task condition: either the relative order, screen locations, or the identity of the pair members. Furthermore, since each relationship required a different kind of effective association, as described above, practice informing relevant kinds of associations to encode the task-critical factor properly was provided before each task.

Within each task section of the experiment, each block began with study trials. At the beginning of each trial, participants viewed a fixation cross in the middle of the screen for 500 ms. As noted above, this was followed by one of the object pictures of the pair, presented either on the top or bottom of the screen, then the second corresponding picture on the opposite part (bottom or top) of the screen, followed by a question mark to indicate that the participant should now make an appropriate association, and finally a blank screen inter-trial interval. To help remind participants which task they should be doing, the object pairs were presented against a color background specific to the condition (temporal – dark blue, spatial – dark green, identity – dark red).

Each encoding block was immediately followed by a test block. As noted, the test trials of each block presented each stimulus only once (and therefore there are only half as many test trials as encoding pairs). Half of the test probes were identical to those seen at encoding, and half were rearranged in a single dimension, as described previously. After the second stimulus of each test pair appeared, participants ranked by a keypress from one to six whether the pair presented was rearranged (1 – definitely rearranged, 2 – fairly sure rearranged, 3 – guess rearranged) or intact (4 – guess intact, 5 – fairly sure intact, 6 – definitely intact). Test pair order was randomized within the block across participants. A 5-minute rest break was given after each task.

Pilot testing indicated that, all other things being equal, the temporal task was the most challenging, followed by the spatial task, with the identity condition task being easiest. In order to identify brain substrates and time courses of processes required for these types of memory, it is important that task difficulty be comparable across task conditions. Therefore, we engaged in extensive iterative pilot testing with the participation of over 100 young adult volunteers who did not participate in the main experiment, in an attempt to balance the difficulty of the tasks using various structural adjustments. The upshot of that adjustment process was that we designed the experiment such that the three experimental tasks were executed in a fixed order, with the easiest task given at the end, when exhaustion and interference are greatest: the temporal task first, then the spatial task, then the identity task. Additionally, each of the tasks employed different numbers of blocks, with different numbers of stimulus pairs in each block. Consequently, the temporal order task had 8 encoding pairs and 4 retrieval trials in each of 4 blocks, the spatial relations task had 16 encoding pairs and 8 retrieval trials in each of 2 blocks, and the associative identity task used a single block of 32 encoding and 16 retrieval trials. In addition, each of the tasks employed different numbers of encoding repetitions. In the identity task, each pair was presented once, and in the temporal and spatial tasks, each pair was presented for encoding twice, in two consecutives but randomly varied sequences. Finally, as noted above, the amount of time given for stimulus display and association formation also differed between conditions (Table 1). We note that in practice there continued to be some differences in task difficulty, as detailed below. Seemingly, these differences would have been even more extreme had we not implemented the differential encoding procedures. We note that we chose to display stimuli above and below fixation, rather than to the right and left of that point, for the benefit of comparison with parallel studies being conducted with the participation of stroke patients who might have hemispatial visual neglect. The entire experiment was presented on a computer running E-Prime 2.0 experimental software (Psychology Software Tools, Pittsburgh, PA, USA).

Regarding the paradigm choice, we should explain that we did not include completely new pairs in the experiment for both fundamental and for logistical reasons. On the fundamental level, trials with item identity novelty (the classic old/new effects case) would have only provided a comparison with the associative identity condition, but not with the critical conditions of temporal and spatial associations. While in principle it might have been nice to have that data as well, just to obtain the basic number of trials in each condition, two recording sessions lasting more than an hour were required. Adding an additional session would, in our estimation, have led to serious participant fatigue which would have confounded the results.

#### 2.4 Electrophysiological Recording Parameters and Data Processing

EEG was recorded throughout the experiment. In the current study, only retrieval-phase data was analyzed.

#### 2.4.1 EEG Recordings

The EEG data was recorded using the Active II system (BioSemi, Amsterdam, The Netherlands) from 64 electrodes mounted in an elastic cap according to the extended 10–20 system. EOG data was recorded using four additional external electrodes, located above and below the right eye and on the outer canthi of both eyes. Additionally, one electrode was placed on the tip of the nose, and two electrodes were placed over the left and right mastoid bones, for reference purposes. The ground function during recording was provided by common mode signal and direct right leg electrodes forming a feedback loop, placed over parieto-occipital scalp. The on-line filter settings of the EEG amplifiers were 0.16–100 Hz. Both EEG and EOG were continuously sampled at 512 Hz and stored for off-line analysis.

#### 2.4.1.1. Preprocessing

Using the Brain Vision Analyzer 2.2 software, stimulus-locked ERPs were segmented into epochs starting 500 msec before the beginning of recognition cue presentation and for 4500 ms afterward. EEG and EOG channels were then referenced to the average of the left and right mastoid channels, band-pass filtered, with an off-line cutoff of 0.5–30 Hz, and baseline-adjusted by subtracting the mean amplitude of the pre-stimulus period (200 ms) of each trial from all the data points in the segment. Independent component analysis was employed to remove heart, eye movements, and blink artifacts. Additional trials containing electrode artifacts and muscle artifacts were rejected visually. Channels depicting drifts and other artifacts in individual trials were replaced with interpolated data from adjacent electrodes.

### 2.5 Statistical analysis

#### 2.5.1 Behavioral analyses

Accuracy level and reaction time (RT; for correct responses) were used as dependent behavioral measures, on which repeated measures ANOVAs were conducted, with associative type (temporal, spatial and identity relations) and associative status (intact, recombined) as repeated factors. Significant effects and interactions were further decomposed using pairwise comparisons and paired t-tests. Here and in all other analyses, Holm–Bonferroni correction was applied to account for inflated type I error due to multiple comparisons. Calculations were executed using the SPSS 25 statistics program (IBM Corp., Armonk, NY, USA).

#### 2.5.2 ERP data segmentation

ERP waveforms were computed for the six retrieval conditions mentioned above (i.e., temporal-intact, temporal-rearranged, spatial-intact, spatial-rearranged and identity-intact, identity-rearranged). For data segmentation, we used 9 representative electrodes covering left anterior (F3), mid anterior (Fz), right anterior (F4), left central (C3), mid central (Cz), right central (C4), left posterior (P3), mid-posterior (Pz) and right posterior (P4) locations (Bridger et al., 2017). Visual inspection of the averaged waveforms revealed a clear peak response during the 300-500 ms window, paralleling the classic FN400 window for recognition memory effects reported in many studies (e.g., Bader et al., 2010; Greve et al., 2007; Kriukova et al., 2013; Opitz, 2010; Wiegand, Bader, & Mecklinger, 2010). The following time window exhibits additional differential development of responses across conditions, and then an inflection point at ∼800 ms before an additional time period of divergence. This may be seen clearly in Figure 3, which portrays the wave forms for each condition separately from three midline electrodes (Fz, Cz, Pz; the waveforms of all nine electrodes contributing to the analysis are presented in Supplementary Figure 1). We therefore separately analyzed the responses for the 500-800 ms and 800-1000 ms time windows. The 500-800 ms time window generally corresponds to the late posterior component of recognition memory effects (e.g., Bader et al., 2010; Greve et al., 2007; Kriukova et al., 2013; Opitz, 2010; Wiegand, Bader, & Mecklinger, 2010). The third time window (800-1000 ms) corresponds to the later part of the range of effect latencies found in ERP recognition studies (reviewed by Mecklinger, 2000; Rugg & Curran, 2007; Wilding & Ranganath, 2011), possibly due to increased demands posed by the retrieval of complex associative information, in contrast to more common item recognition paradigms, most of which have used verbal stimuli (Rugg & Curran, 2007; Tibon et al., 2014). Since the earliest mean manual RTs were at approximately 1250 ms, we did not examine the period after 1000 ms, to exclude the impact of motor preparation on retrieval related activity.

**Figure 2.**
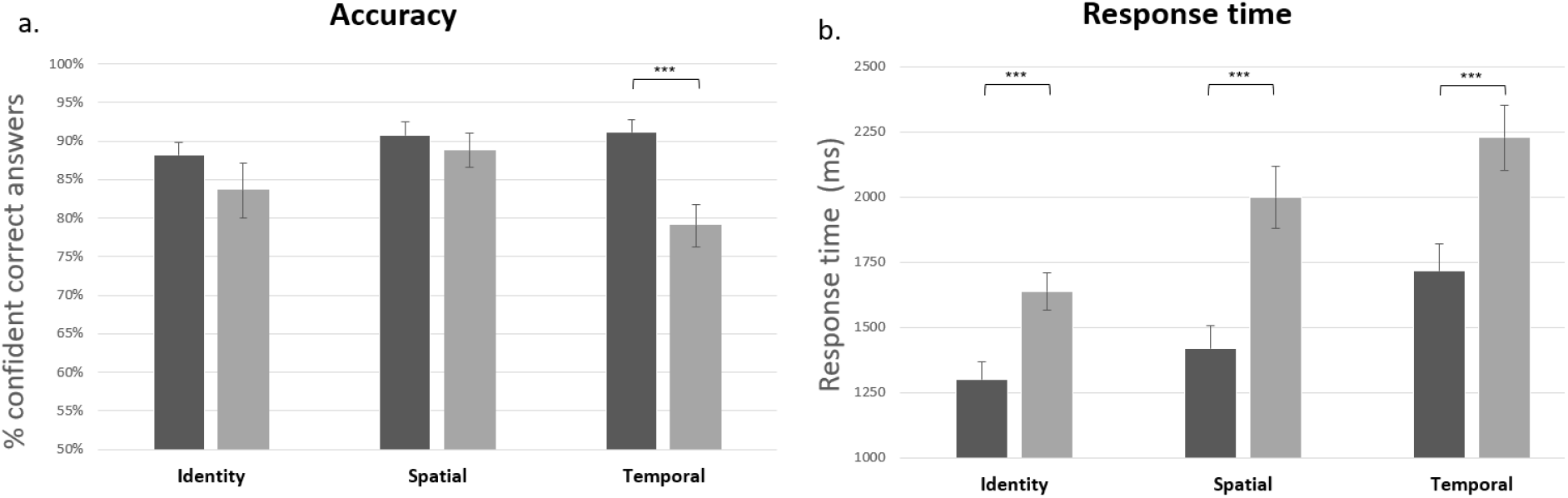
(a) Percent confident correct answers (excluding guesses) for intact (dark color) and recombined (light color) pairs in each associative type (b) Response times (ms) for intact (dark color) and recombined (light color) pairs in each associative type. Error bars indicate SEM. As indicated in Table 2, for both accuracy and response time, main effects of associative type, status, and the interactions between them were significant.

**Figure 3.**
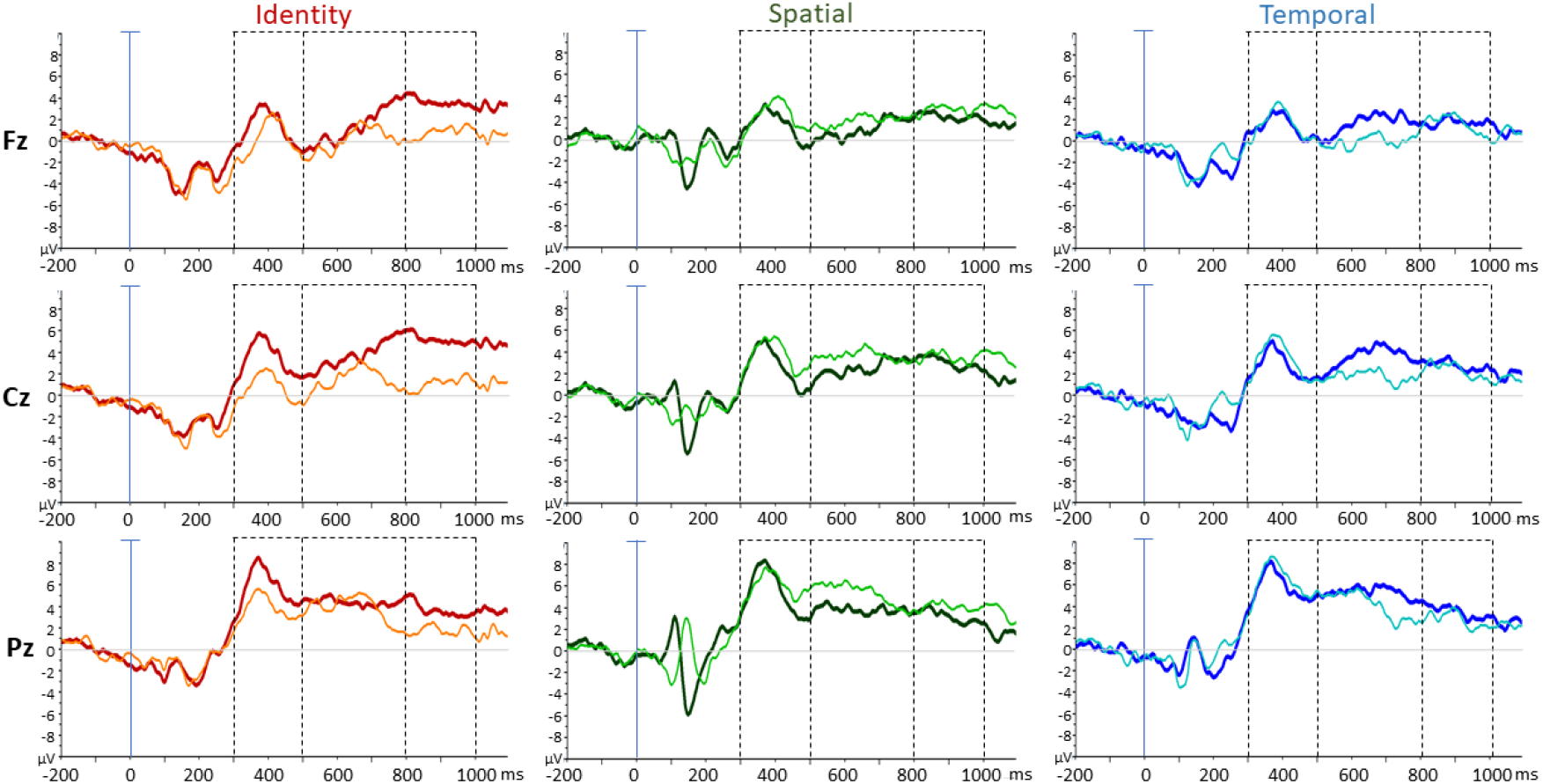
Averaged ERP waveforms elicited by correct recognition of intact (Bold lines) and rearranged stimulus pairs of Temporal, Spatial and Identity relations. Data are shown for the three of nine electrodes used in all statistical analyses. Dashed lines indicate the three-time windows used for statistical analyses.

#### 2.5.3 Mixed-effects models ERP analyses

To analyze the ERP data, we used a linear mixed-effects models approach that was performed separately for each time window. This analysis takes subject-specific variability into account in modeling effects, and can accommodate the repeated measures study design. Such models can be considered a generalization of ANOVA, but use maximum likelihood estimation instead of sum of squares decomposition. An advantage of such an approach over the standard repeated measures ANOVA is that mixed-effects models are better suited for complex designs (e.g., Bagiella, Sloan, & Heitjan, 2000; see also our previous reports, in which we employed a similar approach: Tibon & Levy, 2014a, 2014b; Tibon et al., 2014). Moreover, such an approach is particularly recommended for unbalanced data (Tibon & Levy, 2015), as in the current case in which the number of trials in each condition varied due to differences in accuracy rates between conditions (see Figure 1). Inter-individual differences in EEG amplitude dynamics were modeled as a random intercept, which represents an individual ‘‘baseline,’’ in addition to being affected by the fixed factors. In this mode of analysis, each observation serves as an element of analysis to be modeled; degrees of freedom represent the number of observations and not the number of participants, as is customary in grand average ANOVAs. These parameters result in increased degrees of freedom compared to traditional designs. Although at first glance this might appear to be an overly liberal approach, in this approach large intra-subject variance is not tempered by averaging within participants, which limits the number of effects that emerge as significant. Furthermore, effects that do emerge from the statistical analyses are reflected by robust differences in mean amplitudes. The fixed part of the model includes the associative type factor (temporal, spatial and identity relations), the associative status factor (intact, recombined), and two scalp location factors: anteriority (anterior, central, and posterior) and laterality (left, midline, and right). These scalp locations were represented by the nine representative electrodes mentioned above (F3, Fz, F4, C3, Cz, C4, P3, Pz, P4), as in our previous research and several other prior studies (e.g., Bridger et al., 2017; Czernochowski et al., 2005; Messmer et al., 2020; Stahl et al., 2010). The fixed part of the model further included all possible interactions between these four fixed factors. Model parameters were estimated with the nlme package of the software R (Pinheiro, Bates, DebRoy, Sarkar, & the R Core team, 2007, freely available at http://www.R-project.org).

## 3. Results

We began our analyses with an examination of the experimental design by comparing the average accuracy rates between the two sessions over the three task conditions. We conducted a repeated-measures ANOVA on the dependent measure of percent correct responses, with a within-subject factors of associative type (identity, spatial, temporal) and session time (first session, second session). This analysis revealed a main effect of associative type, *F*(2, 56) = 4.24, *p* = .02, partial η^2^ = .16, but not of session time, *F*(1, 22) = 1.43, *p* = .24, partial η^2^ = .01, nor of the interaction between them, *F*(2, 44) < 1.0. We therefore collapsed the data of the two sessions for further analyses.

We then examined the manipulation employed to equate the difficulty of identity, spatial, and temporal associative memory tasks as detailed in the Methods, by comparing the average accuracy rates across the three task conditions (Identity 85.9%, Spatial 89.8%, and Temporal 85.1%, respectively). These accuracy rates differ significantly, *F*(2, 56) = 3.78, *p* = .03, partial η^2^ = .12, due to the low accuracy rate of the rearranged condition on the temporal relation task (Figure 2a). However, accuracy rates of the intact pairs did not differ significantly across the three task conditions, *F*(2, 56) = 1.30, *p* = .28, partial η^2^ = .04. This indicates that the structural manipulation designed to reduce inter-task difficulty differences was partially successful. However, considering these results, findings about the ERP correlates of responses to the rearranged condition of the temporal relations task were interpretated cautiously.

### 3.1 Behavioral results

Repeated-measures ANOVA was conducted on the dependent measure of percent certain and fairly sure responses (i.e., excluding guess responses), with within-subject factors of associative type (identity, spatial, temporal) and associative status (intact, rearranged). These data are portrayed in Figure 2a.

**Table 2.**
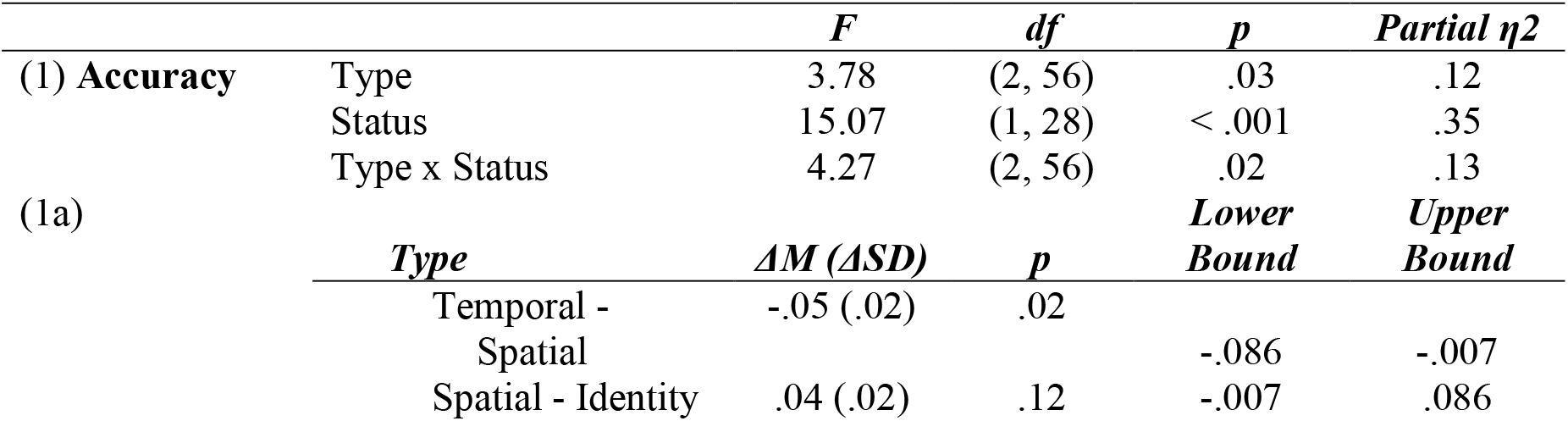
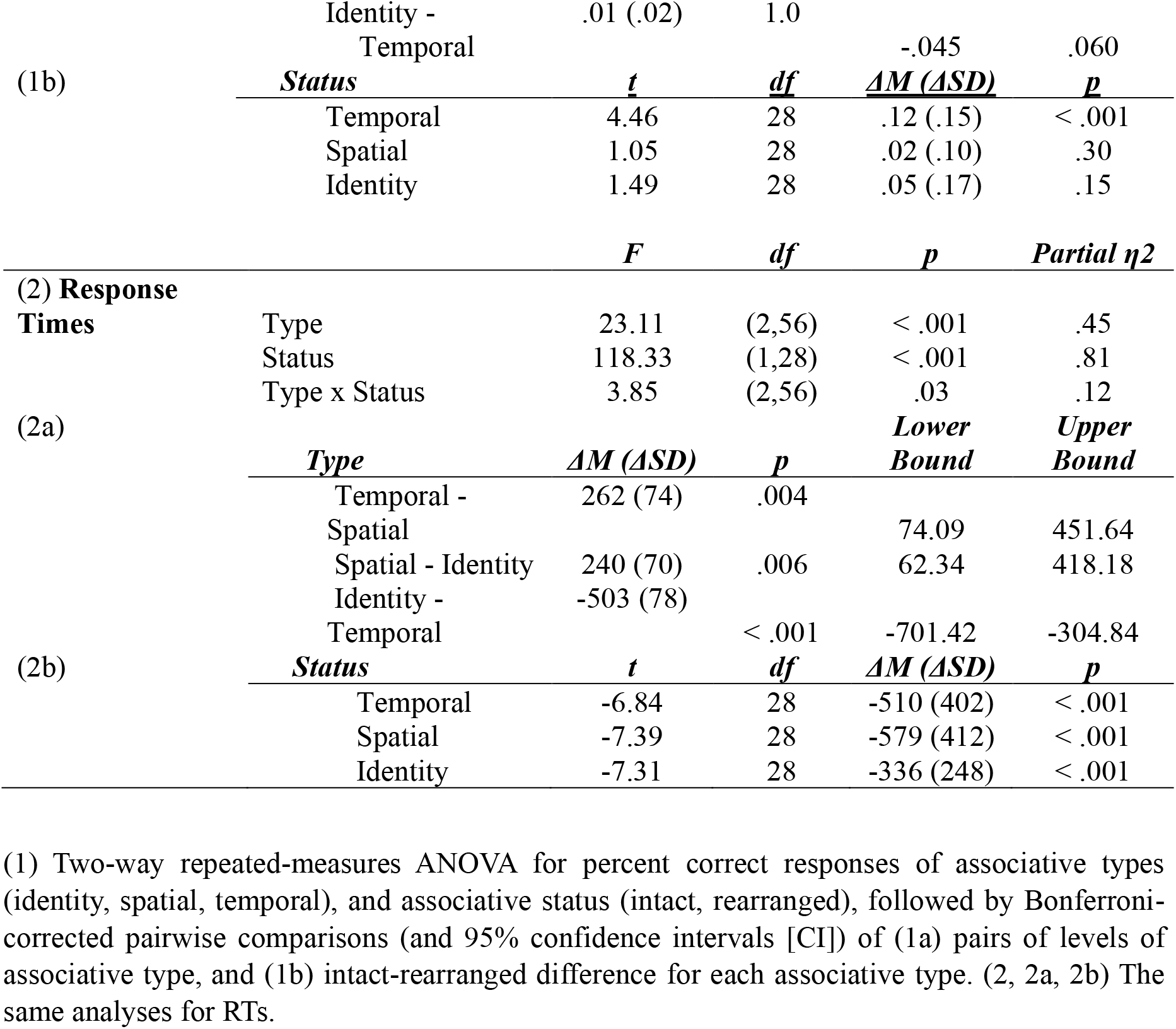
Analysis of variance results for accuracy and response time.

This analysis (Table 2.1) revealed main effects of associative type and associative status, and an interaction between these factors. Bonferroni-corrected pairwise comparisons revealed that across associative status, discrimination of spatial relations was better than discrimination of temporal relations, but the discrimination of identity relations did not differ from spatial or temporal relations. In addition, across all three associative types, participants had relatively greater difficulty in correctly identifying rearranged pairs than intact pairs. However, examination of the interaction between associative status and associative type revealed that identifying pairs as rearranged was significantly more challenging than endorsing intact pairs only for temporal relations. In other words, this data indicates greatest difficulty in correctly identifying temporally rearranged pairs.

We conducted a two-way ANOVA of participant response times for correct responses as the dependent variable, with associative type (identity, spatial, temporal) and associative status (intact, rearranged) as within-subjects factors. Those data are presented in Figure 2b. Analysis of those data (Table 2.2) revealed main effects of associative type and associative status, and significant interactions between these effects. Bonferroni-corrected pairwise comparisons revealed that across associative status, discrimination of spatial relations was faster than discrimination of temporal relations, and identity relations was faster than both spatial and temporal relations. Across all associative types, participants responded faster to intact pairs than rearranged pairs.

### 3.2 ERP results

The first stage of analysis of the recordings involved determining the number of artifact-free EEG segments for correct answer trials in each condition for each participant. That reveled the following distribution: Identity intact -*M* = 11.86, *SD* = 2.85; Identity rearranged -*M* = 11.48, *SD* = 3.45; Temporal intact -*M* = 11.24, *SD* = 4.07; Temporal rearranged -*M* = 8.48, *SD* = 3.60; Spatial intact -*M* = 12.03, *SD* = 3.34; Spatial rearranged -*M* = 10.82, *SD* = 3.74. Since this is a low number of trials for standard grand average ERP analysis, and because of differences in trial numbers between participants, we made planned use of Mixed-effects Models analysis, which we have explained to be the appropriate approach to such data (Tibon & Levy, 2015).

As mentioned above regarding the considerations for data segmentation, the choice of time windows of interest for examination was informed by previous studies and preliminary visual inspection of the overall patterns of responses across conditions. We examined three ERP response segments: a peak response during the 300-500 ms window, paralleling the classic FN400 recognition memory effects, and two following epochs distinguished by an inflection point at ∼800 ms before an additional time period of divergence, yielding time windows of 500-800 ms (paralleling the canonical Late Positive Component [LPC] window) and 800-1000 ms (Figure 3; see also Supplementary Figure 1).

Mean amplitudes in each condition for each task and time window are portrayed in Figure 4, reflecting the data subjected to analysis reported in Table 3. For all three time windows, the mixed-effects model analysis revealed significant main effects of associative type and of anteriority, and significant interactions between associative type and status. In the last two time windows (500-800 ms and 800-1000 ms), there was also a main effect of associative status (Table 3). In the second time window (500-800 ms) there was a significant interaction between associative type and anteriority. To decompose these interactions, for each time window, we collapsed over the laterality factor, which was not significant and did not interact with associative status, and examined effects of associative type and status.

**Table 3.**
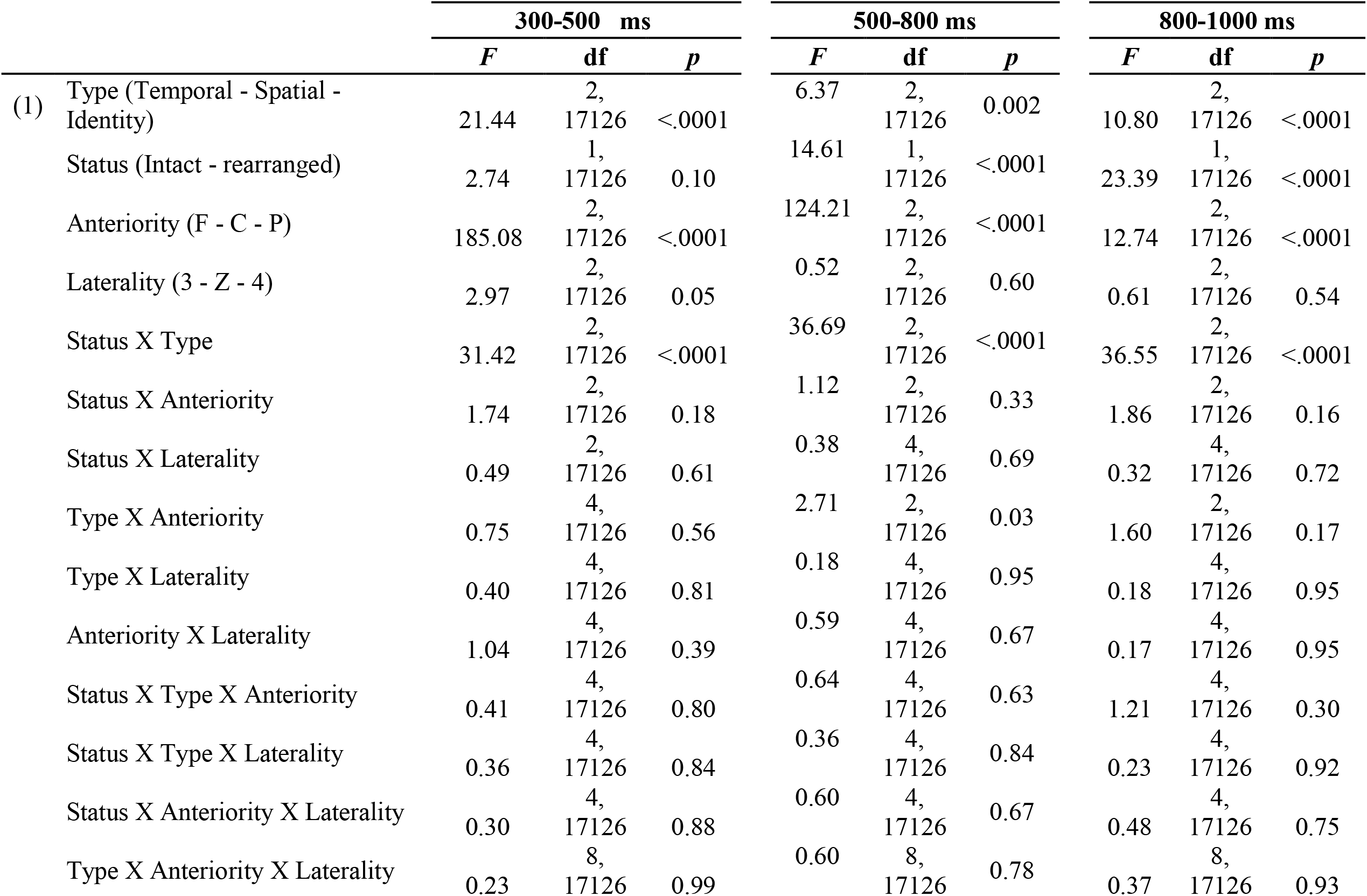
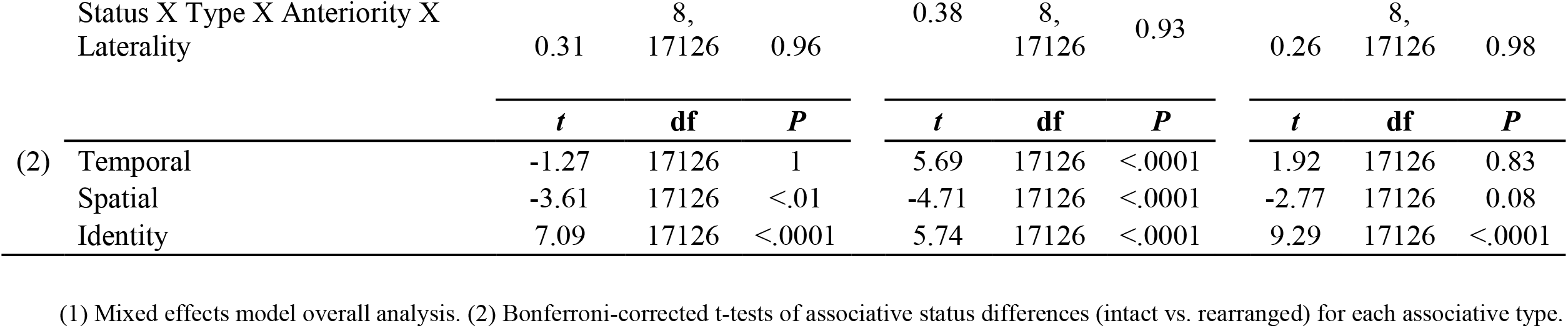
Mean EEG Amplitude - Mixed effects model analysis

**Figure 4.**
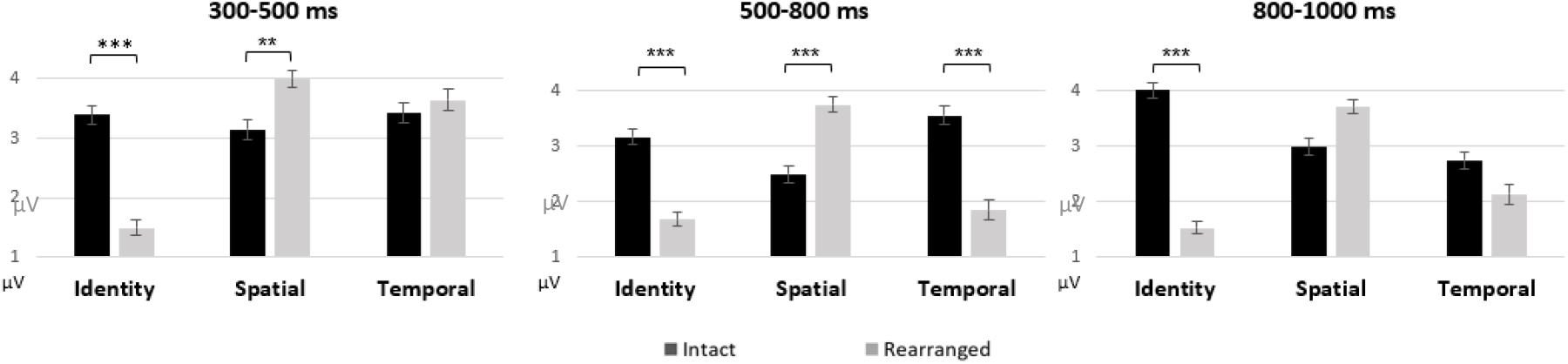
Mean EEG amplitudes for confident correct answers (excluding guesses) in each time window for intact (dark color) and recombined (light color) pairs in each associative type. Error bars indicate SEM.

For the first time window (300-500 ms), this analysis yielded significant main effects of associative type and anteriority, and an interaction between type and status (Table 3). However, anteriority did not interact with associative type in any configuration, and was therefore not included in subsequent analyses. We conducted Bonferroni-corrected t-test comparisons between mean amplitudes associated with intact and rearranged pairs. This analysis revealed significant status differences between intact and rearranged pairs for identity and spatial judgments. However, as is apparent in Figure 4, these differences exhibited different polarities in each associative type: while in identity judgments, ERP deflections were more negative-going for rearranged than the intact pairs, the opposite was true for spatial judgments.

For the second time window (500-800 ms) analysis revealed significant main effects of associative type, status and anteriority, and an interaction between associative type and status (Table 3). Although anteriority did interact with associative type, the anteriority X type X status interaction was not significant, and therefore anteriority was again not included in subsequent analysis. We decomposed the interaction using Bonferroni-corrected t-tests, which revealed a significant difference between responses to intact and rearranged pairs for identity, spatial and temporal relations judgments. However, as is once again apparent from Figure 4, these differences exhibited different polarities in these associative types: while in spatial judgments, ERP deflections were more positive-going for rearranged than for intact pairs, the opposite was true for identity and temporal judgments.

For the third time window (800-1000 ms), we found significant main effects of associative type, status, and anteriority, and an interaction between associative type and status (Table 3). As for the first time window of interest, anteriority did not interact with associative type in any configuration and was therefore not included in subsequent analysis. We decomposed the interaction using Bonferroni-corrected t-tests, which revealed a significant difference between responses to intact and rearranged pairs for identity relations but not for spatial nor for temporal relations judgments.

While planned analyses did not reveal significant interactions between laterality or anteriority and the factors of interest, some additional perspective on the electrophysiological responses associated with discrimination between intact and rearranged pairs of each associative type is provided by the scalp maps of Figure 5, which portray the mean amplitudes of the intact-rearranged voltage differences across the entire scalp for which there was coverage using the 64 electrodes available. Asterisks indicate time windows in which there were significant differences between intact and rearranged status for each type of memory in each time window. This display provides visualization of the results of the statistical analyses that indicated that there were no main or interactive expressions of laterality differences, and no critical interactions between anteriority and task type or associative status. Thus, for these kinds of complex associative recognition of identity, temporal and spatial relations, seemingly widespread mnemonic substrates are recruited, which do not lend themselves to regional limitations of the type associated with right lateralization and frontal displacement of the FN400 component associated with item novelty detection or the left lateralization of the late positive component associated with recollective processes of single-item recognition, which have been documented primarily for verbal materials (see studies listed in Tibon et al., 2014).

**Figure 5.**
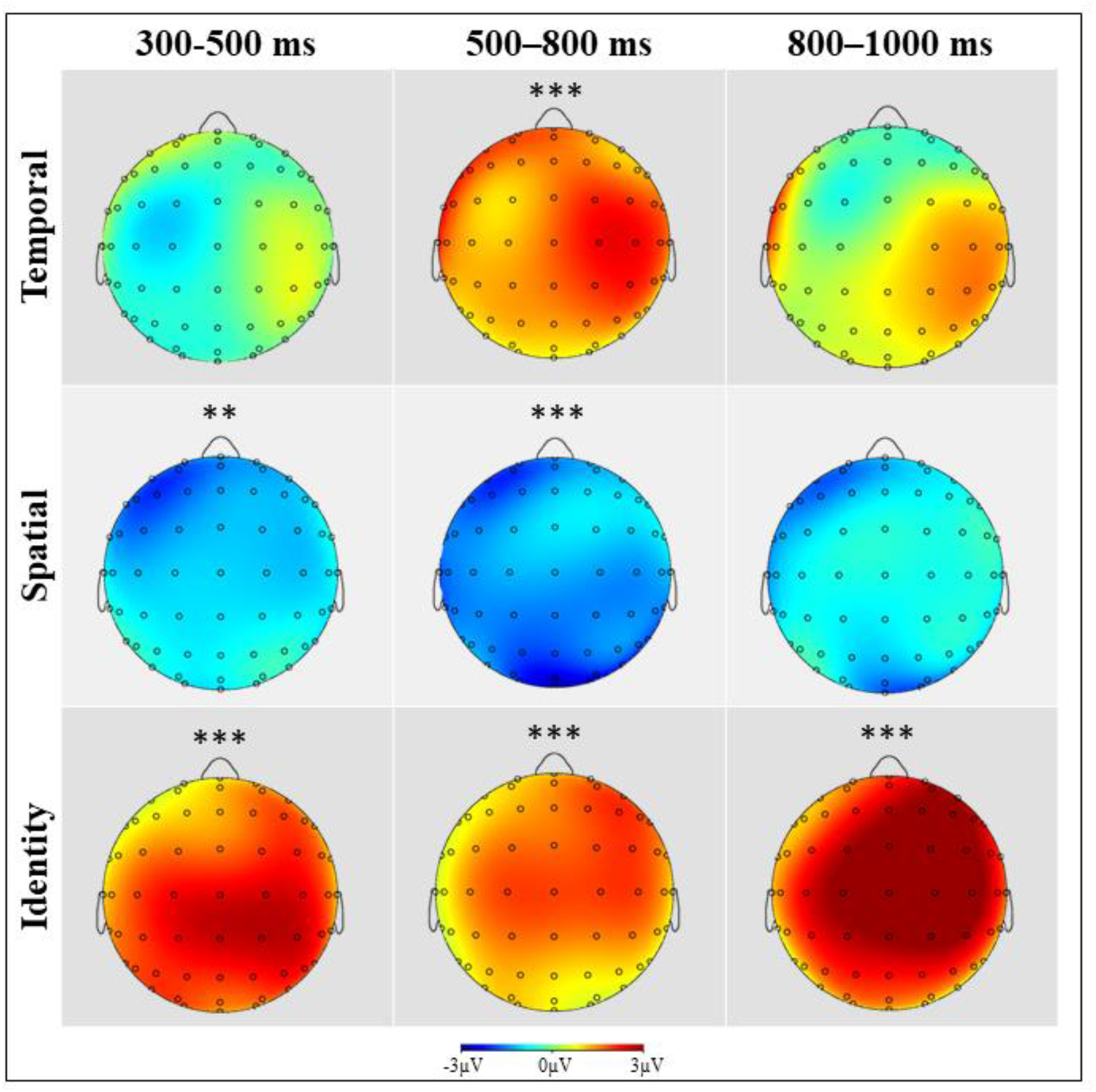
Scalp maps of the mean voltage differences between intact and rearranged pairs for each task type and each time window. Asterisks indicate significant associative status differences in the analyses executed for 9 representative electrodes as reported in Table 3.

## 4. Discussion

In the current study, three minimal-pair associative recognition tasks for temporal, spatial and identity relations were employed to explore whether these three association types require differential retrieval processes, as indexed by their putative electrophysiological signatures. We found that ERP patterns during successful recognition of temporal, spatial and identity relations differed by the status of association. Specifically, for the identity relations condition, the rearranged pairs exhibited a more negative-going deflection than intact pairs in all three time windows. For the spatial relations condition, in the early and the second retrieval stage (300-800 ms) there was a more positive-going deflection for rearranged than for intact pairs, that dissipated in the third epoch (800-1000 ms). In the temporal relations condition, in the second retrieval stage (500-800 ms) there was a more negative-going deflection for rearranged than for intact pairs that dissipated in the final epoch (800-1000 ms).

What types of retrieval processes might be reflected in these dynamic EEG response trajectories? In identity relations judgments, it is possible that the early (300-500 ms) more negative-going deflection evoked by rearranged pairs reflects discrimination between associative novelty and associative familiarity. Although associative memory is generally dependent on recollective processes, in some of our earlier research we found similar early divergences between intact and rearranged pairs of object pictures (Tibon, Ben-Zvi & Levy, 2014). We speculated that bounded presentation of visual objects in same perceptual modality might enable a form of unitization that can allow memory judgments to recruit feelings of associative familiarity or novelty, contributing to a decision that the presented pair is intact or rearranged. We labeled that kind of retrieval ‘direct ecphory’ – a type of retrieval process initiated when a cue enables direct access to the target, and therefore does not require recollective reference to the encoding context. It should be noted that in the aforementioned research, stimuli were presented simultaneously, which might arguably be required in order to create a unitized associative representation that can be accessed without recollective processes. However, in the current study, the assignment of stimuli to discrete locations on screen (rather than appearing in the same location, which could cause masking) might have enabled participants to construct a unitized representation at encoding. That engram could be compared with a similarly unitized mental image of the test probes to engender feelings of familiarity or novelty. While the divergence between the intact and rearranged pairs was significant across all three time windows, examination of the waveforms reveals that after a mid-latency (500-700 ms) transitional state, there once again emerged a late (∼700-1000 ms) status divergence, with more positive amplitude being associated with the intact pairs. This later divergence might be related to the common Late Positive Component often said to reflect recollective processes (Curran & Rugg, 2007), which might have found expression in the ERP patterns later for associative discrimination than for the usual findings for item recognition. Alternatively, since response times in the identity condition were relatively fast, this effect might reflect post-ecphoric evaluation or monitoring processes.

Following this reasoning, the EEG correlates of temporal associative recognition exhibit a more canonical recollective format. There is no divergence between intact and rearranged pairs in the earliest time window. In contrast, we see a divergence reminiscent of the Late Positive Component in the standard time window of 500-800 ms, indicating that recollective judgments might be the key standard of temporal associative discrimination. The divergences are not observed in the final time window, possibly because recollection might not require the kind of monitoring evaluation related to familiarity. Temporal relations recognition may be a profoundly different process from retrieval of spatial and identity relations, as the temporal task could not be accomplished by retrieving just one visual mental image, but require the participant to retrieve the two serial components of the episode (Kwok et al., 2012). Interestingly, according to chronological organization theory (Friedman, 1993), retrieving temporal relations can be done by retrieving the time position of one of the two objects in the episode, and then scanning through the rest of the episode looking for the second object (i.e., serial temporal search). The character of the EEG correlates of temporal relations task might reflect this serial temporal search that involve unique cognitive functions.

An alternative explanation of the profile of effects in the temporal relations condition should be noted. The manipulation described above that reduced task-difficulty differences between the conditions resulted in the temporal relations condition being based on 8 encoding trials and 4 retrieval trials. While this still represents a considerable amount of temporal and associative information, it is conceivable that working memory retention might have contributed more to retrieval in that condition than in the other conditions which employed longer block length.

The spatial relations condition exhibits a very different pattern of activity than the other two conditions. Across time windows, it was characterized by greater positivity in response to the rearranged pairs (the opposite polarity of what is found in the other conditions and in canonical old/new effects), most prominently in a time window 400-800 ms after presentation of the second probe. Such a pattern is not attested in earlier studies, and therefore it is challenging to interpret it with any certainty. We note, however that relative polarity of components have been reported to be strongly influenced by task context. For example, Leynes and Upadhyay (2022) have noted that picture fluency (e.g., Bruett & Leynes, 2015; Voss & Paller, 2010) or perceptual manipulations of word stimuli (e.g., Leynes & Zish, 2012) have been associated with an ERP in which old items are more negative than new – i.e., exhibiting the same polarity reversal that we see for spatial condition. We might therefore speculate, based on the time window of the effect, that it appears to reflect processes that are predominantly recollective. Indeed, to the best of our knowledge, there is no evidence in the literature of associative familiarity in memory for spatial relations. However, this recollective process appears to begin earlier for spatial than for temporal relations, and extends later. This unusual pattern of results raises the possibility that the type of recollective processing supporting spatial associative memory might be different from those supporting identity or temporal associative memory. This is a possibility that seems worthy or investigation in future studies.

Following that processing, in the spatial relations condition, as we noted for temporal associative judgments, there are no further status condition differences in the final time window. Again, this may be because decisions based solely on recollection may not require post-ecphoric monitoring to the same extent as familiarity-based judgments. Alternatively, since response times for temporal and spatial associations are much longer than those for identity relations, it is possible that a post-retrieval evaluative or monitoring process might occur at some later point that we could not analyze in the current data, due to overlap with motor response preparation, as noted above.

Comparison of the present results with prior EEG studies of associative memory, which might have provided traction on the interpretation of the current results, is not simple for a number of reasons. One reason is that almost all such studies made use of words as memoranda (e.g., Kriukova et al., 2013; Rhodes & Donaldson, 2007, 2008), while the present study used object pictures as memoranda. Furthermore, an important feature of the present study is that in order to examine temporal order memory, the stimuli were always offset in time, at encoding and at retrieval. The associative recognition decisions could only be made upon onset of the second stimulus. To the best of our knowledge, no studies of associative recognition have featured such offsets; Tibon and Levy (2014) used offset at encoding, but examined cued recall processes. Differences in EEG patterns between prior studies and the current experiment might be a factor of that feature. It is possible that simultaneous presentations might have enabled use of associative familiarity signal to make identity and spatial associative recognition judgments, with ensuing early FN400 strengthened (in the case of identity; as it is observed even in the current paradigm) or emerging (in the case of spatial relations).

As noted in the introduction, one of the few studies that have equated cognitive strategies and retrieval performance between spatial versus temporal tasks was performed by Rajah and colleagues (2011). The researchers reported greater brain activity in the left anterior PFC, bilateral VLPFC and right DLPFC during spatial vs. temporal context retrieval. However, most of these PFC regions were also more active during retrieval than during encoding and were more engaged during difficult vs. easy retrieval events. The researchers concluded that PFC contributions to spatial and temporal retrieval may not reflect inherent differences in the cognitive control processes specifically important for retrieving spatial versus temporal context information, rather reflecting this region’s importance in various domain-general cognitive control processes (Stuss & Knight, 2002). Our finding of dynamic differences in EEG responses to intact and rearranged pairs, alongside the absence of laterality differences within each time window, suggest that these regions might contribute to both spatial and temporal associative memory retrieval, but that their contributions might vary over time in a task-dependent manner.

Other studies have proposed stronger domain specificity of retrieval-related processes (Kwok et al., 2012; Kwok & Macaluso, 2015). In these fMRI studies, participants made mnemonic judgments following a short video clip and were asked to choose either the scene that happened earlier in the film (temporal), or the scene with a correct spatial arrangement (spatial), or the scene that had been shown (identity). The researchers reported that the precuneus and the right angular gyrus were associated with temporal-order retrieval, the dorsal fronto-parietal network engaged during spatial-related judgments, and antero-medial temporal along with medial frontal regions activated during object-related retrieval. It is possible that the precuneus and the right angular gyrus activity associated with temporal relations retrieval is represented by the mid-latency positive-going deflection for intact relative to rearranged pairs (500-800 ms), for which we have suggested a recollective basis. This is in consonance with the inclusion of the precuneus and angular gyrus in the putative core recollection network (Rugg & Vilberg, 2013). Similarly, the dorsal fronto-parietal network reported to engage during spatial relations retrieval might be associated with the intact-rearranged deflection differences observed in the first and second time windows. It is also possible that the activation reported in the medial regions during scene recognition (Kwok & Macaluso, 2015) is reflected in our data in the divergence between rearranged and intact pairs in the early stage (300-500 ms), which we have suggested might represent associative familiarity. Of course, many process accounts of the fMRI results are possible, since in univariate hemodynamic analysis, the dynamically differential activation of neuronal populations within a given region that lead to shifting polarities in the EEG signal are not distinguished in a single TR. Employment of intracranial electrode recording, or simultaneous fMRI-EEG, will be required to test this speculation.

## 5. Conclusions

The current study provides important traction on the question of processes underlying associative episodic memory for temporal, spatial, and identity relations. Going beyond our previous findings that healthy aging seems to equally impact all three types of remembering (Hugeri et al., 2022), the clear electrophysiological differences which we report indicate that each type of relationship might dynamically recruit multiple retrieval processes. Additional research providing more information about anatomical substrates of these processes is required. We are currently conducting a neuropsychological lesion effects study using the same experimental paradigm, which, together with hemodynamic imaging and other approaches, will hopefully more fully illuminate this important aspect of how we remember the events of our lives.

## Acknowledgement

The authors wish to thank Tal Gigi for assistance with data collection, and Shir Ben-Zvi Feldman for assistance regarding data analysis. D.A.L. and O.H. are supported by the Israel Science Foundation, Grant 2497/18.

## AUTHOR CONTRIBUTIONS

Ofer Hugeri: Conceptualization; investigation; formal analysis; writing – original draft; writing – review and editing. Eli Vakil: Methodology; supervision; writing – review and editing. Daniel A. Levy: Conceptualization; methodology; project administration; resources; supervision; validation; writing – review and editing.

## Supplementary Material

**Supplementary Table 1.** Randomization display of stimuli for the Temporal, Spatial and Identity relations at each experiment

| Temporal experiment |  |  |  |  |  |  |  |
| --- | --- | --- | --- | --- | --- | --- | --- |
| Encoding |  |  |  | Retrieval |  |  |  |
| Up first | Up second | Down first | Down second | Up first | Up second | Down first | Down second |
|  | pizza | book shelf |  |  | pizza | book shelf |  |
| pizza |  |  | feather |  |  |  |  |
|  | blush brush | screw |  |  |  |  |  |
| blush brush |  |  | gloves |  | blush brush | gloves |  |
|  | toaster | feather |  |  | toaster | feather |  |
|  | cigarette | gloves |  |  |  |  |  |
| toaster |  |  | book shelf |  |  |  |  |
| cigarette |  |  | screw |  | cigarette | screw |  |
| Spatial experiment |  |  |  |  |  |  |  |
| Encoding |  |  |  | Retrieval |  |  |  |
| Up first | Up second | Down first | Down second | Up first | Up second | Down first | Down second |
| pizza |  |  | book shelf | pizza |  |  | book shelf |
|  | feather | pizza |  |  |  |  |  |
| blush brush |  |  | gloves |  | gloves | blush brush |  |
|  | screw | blush brush |  |  |  |  |  |
|  | book shelf | toaster |  |  |  |  |  |
| toaster |  |  | feather | toaster |  |  | feather |
|  | gloves | cigarette |  |  |  |  |  |
| cigarette |  |  | screw |  | screw | cigarette |  |
| Identity experiment |  |  |  |  |  |  |  |
| Encoding |  |  |  | Retrieval |  |  |  |

| Up first | Up second | Down first | Down second | Up first | Up second | Down first | Down second |
| --- | --- | --- | --- | --- | --- | --- | --- |
| pizza |  |  | bookshelf | pizza |  |  | book shelf |
| pizza |  |  | feather |  |  |  |  |
| blush brush |  |  | gloves | blush brush |  |  | feather |
| blush brush |  |  | screw |  |  |  |  |
| toaster |  |  | bookshelf | toaster |  |  | screw |
| toaster |  |  | feather |  |  |  |  |
| cigarette |  |  | gloves | cigarette |  |  | gloves |
| cigarette |  |  | screw |  |  |  |  |

### Supplementary Figures

**Figure S1.**
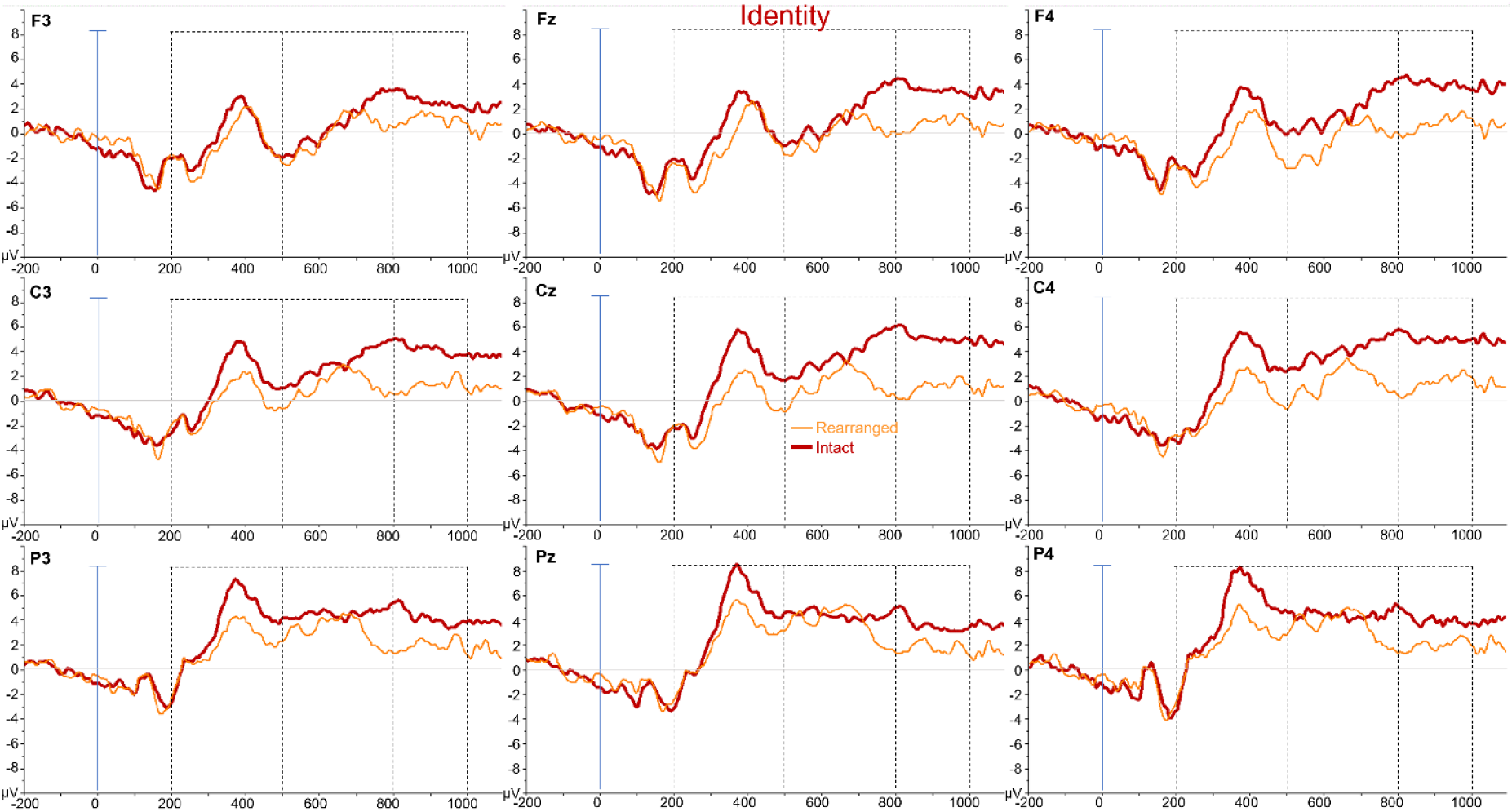

**Figure S2.**
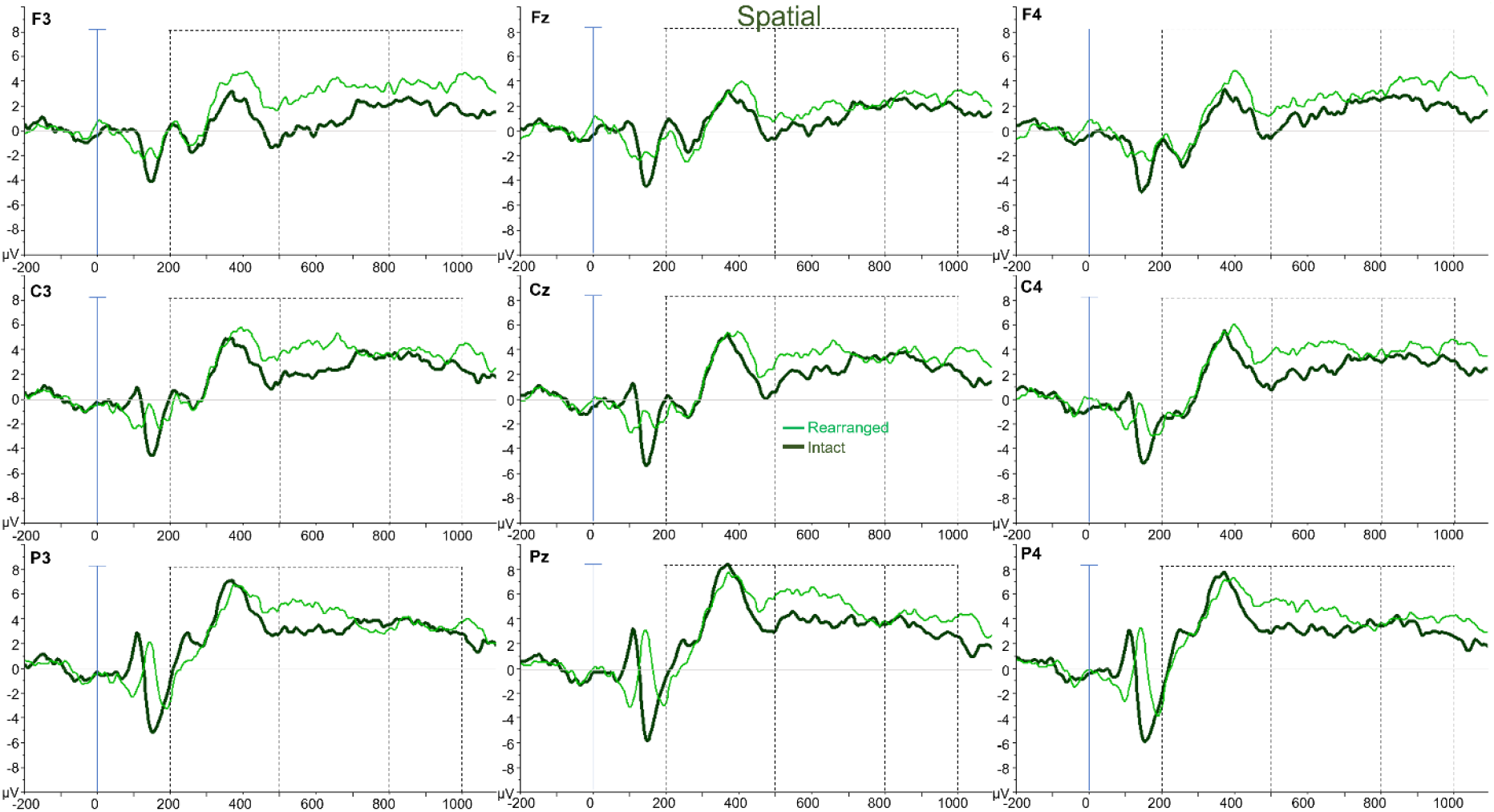

**Figure S3.**
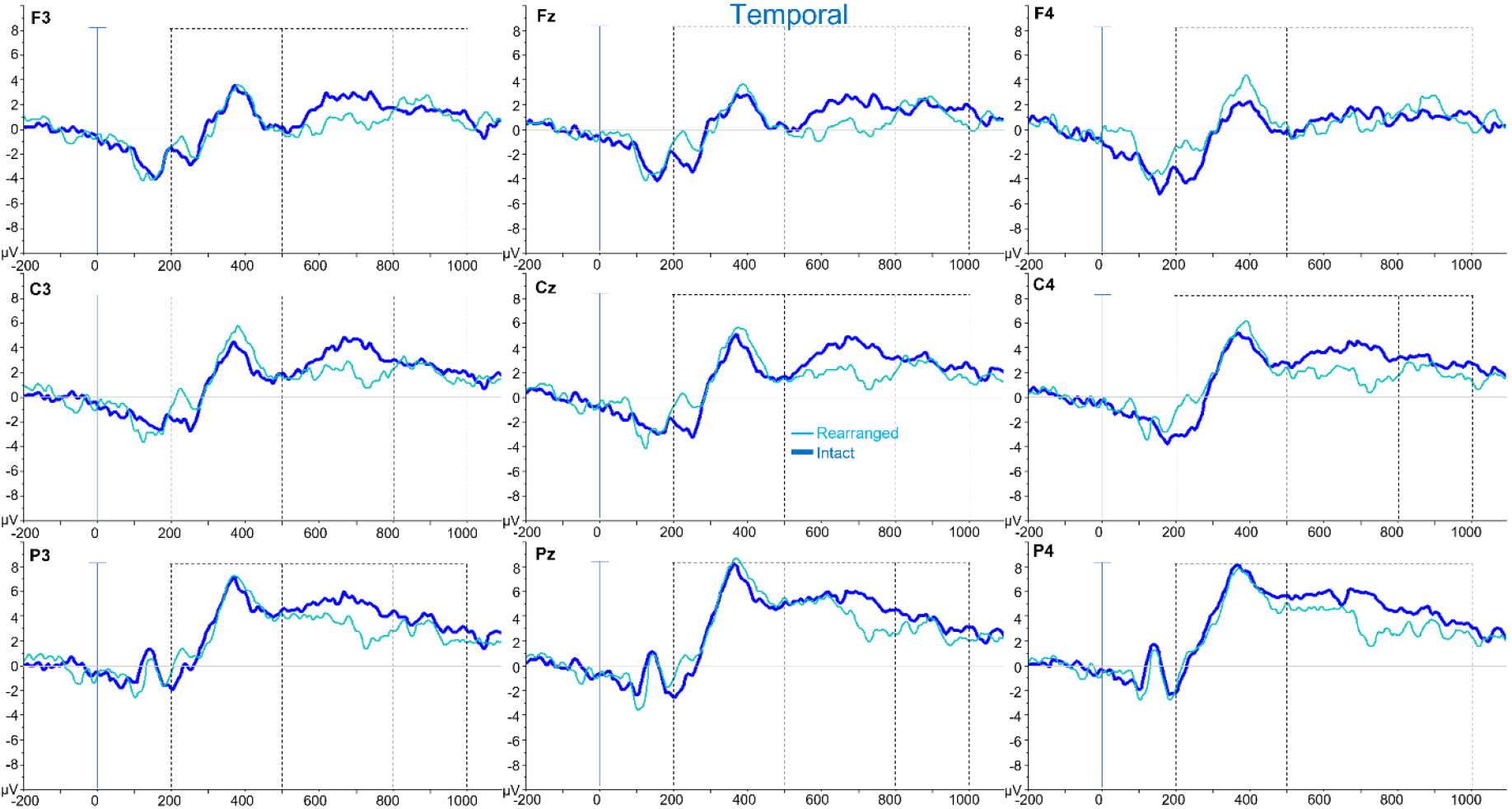
Averaged ERP waveforms elicited by correct recognition of intact (Bold lines) and rearranged stimulus pairs of Identity (Figure S1), Spatial (Figure S2) and Temporal (Figure S3) relations. Data are shown for the nine electrodes used in all statistical analyses. Shadings indicate the three-time windows used for statistical analyses.

